# Virome heterogeneity and connectivity in waterfowl and shorebird communities

**DOI:** 10.1101/528174

**Authors:** Michelle Wille, Mang Shi, Marcel Klaassen, Aeron C. Hurt, Edward C. Holmes

**Affiliations:** WHO Collaborating Centre for Reference and Research on Influenza, at The Peter Doherty Institute for Infection and Immunity, Melbourne, Australia.; Marie Bashir Institute for Infectious Diseases and Biosecurity, Charles Perkins Centre, School of Life and Environmental Sciences and Sydney Medical School, The University of Sydney, Sydney, Australia.; Centre for Integrative Ecology, Deakin University, Geelong, Australia.

**Author notes:** Corresponding author: Michelle Wille, Edward C. Holmes.

**Keywords:** virus, evolution, host - pathogen interactions, influenza A virus, virome, wild birds

## Abstract

Models of host-microbe dynamics typically assume a single-host population infected by a single pathogen. In reality, many hosts form multi-species aggregations and may be infected with an assemblage of pathogens. We used a meta-transcriptomic approach to characterize the viromes of nine avian species in the Anseriformes (ducks) and Charadriiformes (shorebirds). This revealed the presence of 27 viral species, of which 24 were novel, including double-stranded RNA viruses (*Picobirnaviridae* and *Reoviridae*), single-stranded RNA viruses (*Astroviridae, Caliciviridae, Picornaviridae*), a retro-transcribing DNA virus (*Hepadnaviridae*), and a single-stranded DNA virus (*Parvoviridae*). These viruses comprise multi-host generalist viruses and those that are host-specific, indicative of both virome connectivity and heterogeneity. Virome connectivity was apparent in two well described multi-host virus species (avian coronavirus and influenza A virus) and a novel *Rotavirus* species that were shared among some Anseriform species, while heterogeneity was reflected in the absence of viruses shared between Anseriformes and Charadriiformes. Notably, within avian host families there was no significant relationship between either host taxonomy or foraging ecology and virome composition, although Anseriform species positive for influenza A virus harboured more additional viruses than those negative for influenza virus. Overall, we demonstrate complex virome structures across host species that co-exist in multi-species aggregations.

## Introduction

Many hosts are members of multi-species aggregations and may be infected by an assemblage of specialist and/or multi-host generalist infectious agents. Host community diversity is central to pathogen dynamics (1,2), and pathogen species richness, relative abundance, specificity and intra- and inter-species interactions within assemblages likely have complex roles in modulating pathogen levels within populations (1,3-8). A significant limitation in studying viral communities in hosts is that most viral species remained undescribed (9), such that viral ecology across multi-host systems has been limited to “single-virus” dynamics, particularly in vertebrate systems (for example, Influenza A virus [IAV] in avian populations). With the advent of unbiased, bulk ‘meta-transcriptomic’ RNA sequencing we can now explore, in more detail, how viral community structure may be shaped by host-species interactions.

Birds of the orders Anseriformes and Charadriiformes, the central reservoirs for avian viruses such as IAV, avian avulavirus and avian coronavirus (10,11), form multi-host flocks, in which many species may migrate, forage, or roost together (e.g. (12). These flocks may comprise species along a taxonomically related gradient and may utilize similar or different ecological niches in the same environment. For example, in Australia, taxonomically related dabbling Grey Teals (*Anas gracilis*) and Pacific Black Ducks (*Anas superciliosa*) may share the environment with the distantly related filter feeding Pink-eared Duck (*Malacorhynchus membranaceus*). These multi-host flocks form multi-host maintenance communities (6), with consequences for virus ecology, transmission, and virulence (1,13,14).

Studies of the ecology of IAV, the best studied multi-host virus in wild birds, have shown that not all hosts are equal (15,16). In particular, there are marked differences in susceptibility, pathology and the subsequent immune response in taxonomically related species, or distantly related species within similar ecological niches. For example, dramatic differences in viral prevalence exist within the Charadriiformes, such that Ruddy Turnstones (*Arenaria interpres*) may have an IAV prevalence of ∼15%, compared to the negligible IAV prevalence in co-sampled Sanderlings (*Calidris alba*) sampled at Delaware Bay, USA (17). There are also major differences in the pathology of highly pathogenic IAV in Anseriformes both in the field and in experimental infections. Mallards (*Anas platyrhynchos*) infected with the highly pathogenic IAV are thought to move the virus large distances and remain free of clinical signs, while Tufted Ducks (*Aythya fuligula*), in contrast, experience severe mortality (18-20). Following IAV infection, dabbling ducks of the genus *Anas* are believed to develop poor immune memory (21), allowing IAV re-infections throughout their lives, in contrast to swans that have long term immune memory (22) and where re-infection probability is likely very low in adults. These differences are driven by factors encompassing both virus (e.g. virulence, transmission) and host (e.g. receptor availability, immune responses) biology (11).

The goal of this study was to use the analysis of comparative virome structures, particularly virome composition, diversity and abundance, as a means to describe the nature of host-virus interactions beyond the “one-host, one-virus” model. Given their role as hosts for multi-host viruses such as IAV, coronaviruses and avian avulaviruses, we used apparently healthy members of the Anseriformes and Charadriiformes as model avian taxa in these comparisons. In particular, we aimed to (i) reveal the viromes and describe novel viruses in the bird taxa sampled, (ii) determine whether viromes of different host orders have different abundance and viral diversity such that they cluster separately, (iii) determine whether closely taxonomically related and co-occuring avian hosts have viromes that are more homogenous, and (iv) identify the impact of host taxonomy, which we use as a proxy for differences in relevant host traits (such as host physiology, cell receptors), in determining virome structure. Overall, we reveal a combination of virome heterogeneity and connectivity across avian species that are important reservoirs of avian viruses.

## Methods

### Ethics statement

This research was conducted under approval of the Deakin University Animal Ethics Committee (permit numbers A113-2010 and B37-2013). Banding was performed under Australian Bird Banding Scheme permit (banding authority numbers 2915 and 2703). Research permits were approved by the Department of Environment, Land, Water and Planning Victoria (permit numbers 10006663 and 10005726)

### Sample selection

Samples were collected as part of a long-term IAV surveillance study (23,24). Shorebirds were captured using cannon nets at the Western Treatment Plant near Melbourne (37°59′11.62′′S, 113 144°39′38.66′′E) during the same sampling event (n=434). Waterfowl were sampled post-mortem (within 12 hours) following harvest from lakes in south-west Victoria in March 2017 (n=125) (36°58’S 141°05′E) (Table S1). No birds showed any signs of disease.

A combination or oropharyngeal and cloacal samples were collected using a sterile-tipped swab and were placed in viral transport media (VTM, Brain-heart infusion broth containing 2×106 122 U/l penicillin, 0.2 mg/ml streptomycin, 0.5 mg/ml gentamicin, 500 U/ml amphotericin B, Sigma).

### RNA virus discovery

RNA was extracted and libraries constructed as per (25) (Table S1). Paired end sequencing (100bp) of the RNA library was performed on an Illumina platform at the Australian Genome Research Facility (AGRF, Melbourne). Similarly, sequence reads were demultiplexed and contigs assembled and contig abundance calculated as per (25).

All contigs that returned blast hits with paired abundance estimates were filtered to remove plants and invertebrate reads that likely correspond to the host diet, as well as fungal, bacterial and host sequences. The virus list was further filtered to remove viruses with likely invertebrate (26), lower vertebrate (27), plant or bacterial host associations using the Virus-Host database (http://www.genome.jp/virushostdb/) such that only those viruses that grouped within the previously defined vertebrate virus groups are identified as bird associated.

### Virus genome annotation and phylogenetic analysis

Contigs were annotated, and phylogenetic treesinferred as per (25). Briefly, contigs greater than 1000bp in length were inspected using Geneious R10 (Biomatters, New Zealand), and open reading frame corresponding to predicted genome architectures based on the closest reference genomes were interrogated. Reads were subsequently mapped back to viral contigs to identify mis-assembly using bowtie2 (28). Viruses with full length genomes, or incomplete genomes but that possess the full-length RNA-dependant RNA polymerase (RdRp) gene, were used for phylogenetic analysis. Briefly, sequences of the polyprotein or gene encoding for the RdRp were aligned using MAFFT (29), and gaps and ambiguously aligned regions were stripped using trimAL (30). Final alignment lengths for each data set are presented in Table S2. The most appropriate amino acid substitution model was then determined for each data set, and maximum likelihood trees were estimated using PhyML 3.0 (31) with 1000 bootstrap replicates. Novel viral species were identified as those that had <90% RdRp protein identity, or <80% genome identity to previously described viruses. All reads have been deposited in the Short Read Archive (PRJNA505206) and viral sequences described have been deposited in GenBank (MK204384-MK20441, MK213322-MK213337).

### Diversity and abundance across libraries

Relative virus abundance was estimated as the proportion of the total viral reads in each library (excluding rRNA). All ecological measures were calculated using the data set comprising viruses associated with “higher” vertebrates (i.e. birds and mammals), albeit with all retroviruses and retrotransposons removed (hereafter, “avian virus data set”).

Analyses were performed using R v 3.4.0 integrated into RStudio v 1.0.143, and plotted using *ggplot2*. Both the observed virome richness and Shannon effective (i.e. alpha diversity) were calculated for each library at the virus family and genus levels using the Rhea script sets (32), and compared between avian orders using the Kruskal-Wallis rank sum test. Beta diversity was calculated using the Bray Curtis dissimilarity matrix and virome structure was plotted as a function of nonmetric multidimensional scaling (NMDS) ordination and tested using Adonis tests using the *vegan* (33) and *phyloseq* packages (34). To determine whether differences in virome composition can be explained by host phylogeny (35), dendograms of beta diversity were constructed using the bray curtis dissimilarity matrix using the *vegan* package incorporated into the hclust() function. Dendograms representing library abundance and composition at the viral family, genera and species level of taxonomy were also compared to the host phylogeny. For the species level comparison, we used only those presented in Table S3. Two phylogenetic tree congruence metrics were then calculated to compare the match between the virome metric and host phylogeny: the matching cluster Robinson-Foulds tree metric (36) was calculated using the *phangorn* package (37), and the normalized PH85 metric (38) (39) was determined using the *ape* package (40). For both metrics, a distance of 0 indicates complete congruence and 1, incomplete congruence. The phylogenetic relationships among the avian host species were inferred using a maximum likelihood tree of the cytochrome B mitochondrial sequences and accords with those determined previously (41-44). The overall co-phylogenetic analysis was visualized using the *phytools* package (45).

Finally, the relative abundance of virus families between each avian host order (Charadriiformes versus Anseriformes) was assessed using the wilcox test. Subsequently, log2 relative abundance was calculated using *DESeq2* (46) implemented in *phyloseq* (34). Given the large number of genera detected within the *Picornaviridae*, this analysis was repeated at the viral genus level in this case.

## Results

### Substantial undescribed diversity of RNA viruses in wild birds

We characterized the total transcriptome of nine avian species from the Anseriformes and Charadriiformes (Table S1). Within each avian order, bird species were sampled across the same spatial and temporal scales, although members of the Anseriformes and Charadriiformes were sampled at different time points (i.e. years) and locations. This resulted in two bird communities comprising five and four species, respectively (Table S1). RNA sequencing of rRNA depleted libraries resulted in an average of 50,906,440 reads (range 44,178,936-64,149,180) assembled into a mean of 27,108 contigs (range 22,175-496,319) (Table S1). There was no significant correlation between RNA concentration and the number of reads, contigs or abundance demonstrating such that this was not a source of bias (Fig S1). There was a large range in total viral abundance (0.14-10.67% viral reads) and putative avian viral abundance (0.00083-0.327% viral reads) in each library (Table S1, Fig 1). In additional to avian viral reads, libraries had numerous reads matching arthropod viruses and retroviruses (Fig 1). Although these retroviruses are likely bird associated, the challenge of differentiating between endogenous and exogenous retroviruses meant that they were excluded from the analysis, as were those viruses most likely associated with arthropods, plants and bacteria.

**Figure 1.**
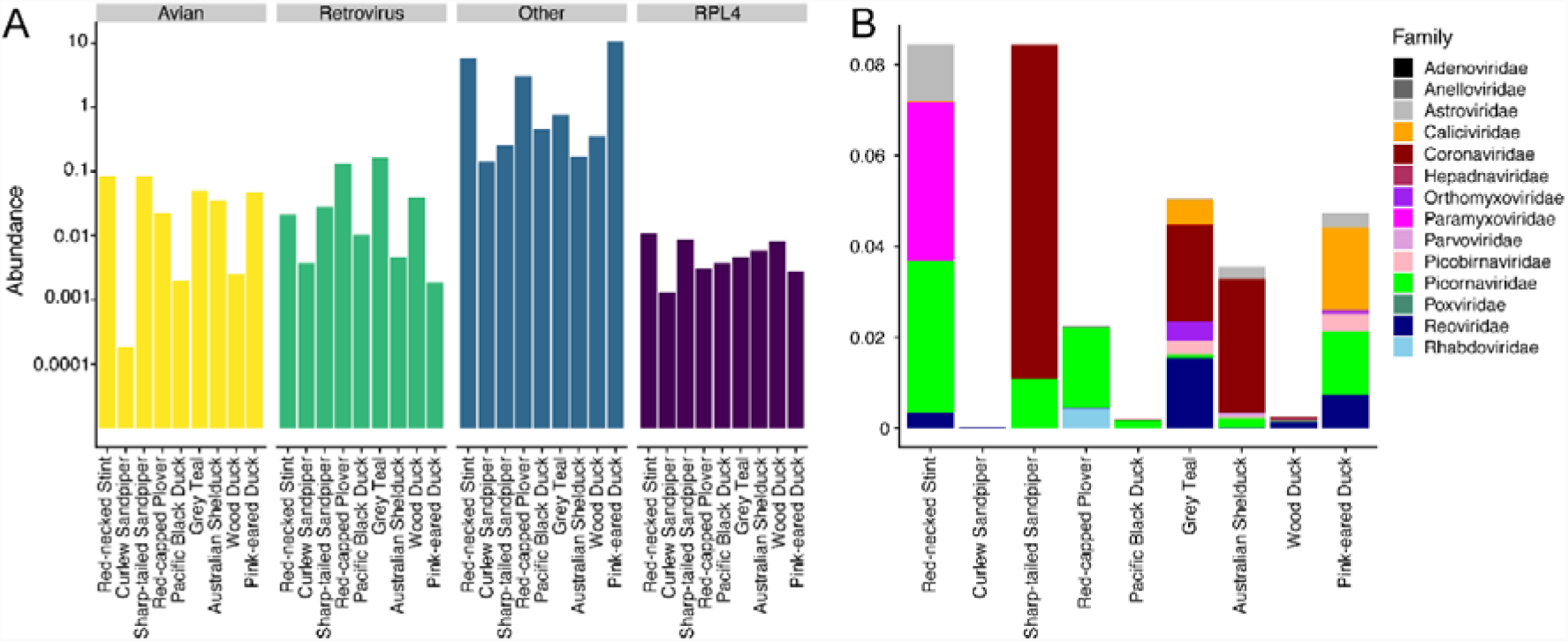
Overview of viral contigs identified in this study. (A) Host association of viral contigs identified in this study: avian, retroviruses and retrotransposons, all other hosts including lower vertebrate, invertebrate, plant, bacterial hosts, and host reference gene ribosomal protein L4 (RPL4). (B) Abundance and alpha diversity of avian viral families identified in each library. Relative abundance and alpha diversity calculations are presented in Fig S15. Abundance and alpha diversity of viral genera and species is presented in Fig S16 and S17, respectively.

A total of 24 of the 27 viruses identified in this study likely represent novel avian viral species (Table S3, Fig 2, Fig S2). Novel species were identified in the double-stranded RNA viruses (*Picobirnaviridae* and *Reoviridae*, genus Rotavirus), positive-sense single-stranded (ss) RNA viruses (*Astroviridae, Caliciviridae, Picornaviridae* genus Megrivirus, Gallivirus, and unassigned genera), and both retro-transcribing (*Avihepadnaviridae*) and single-stranded DNA viruses (*Parvoviridae*).

**Figure 2.**
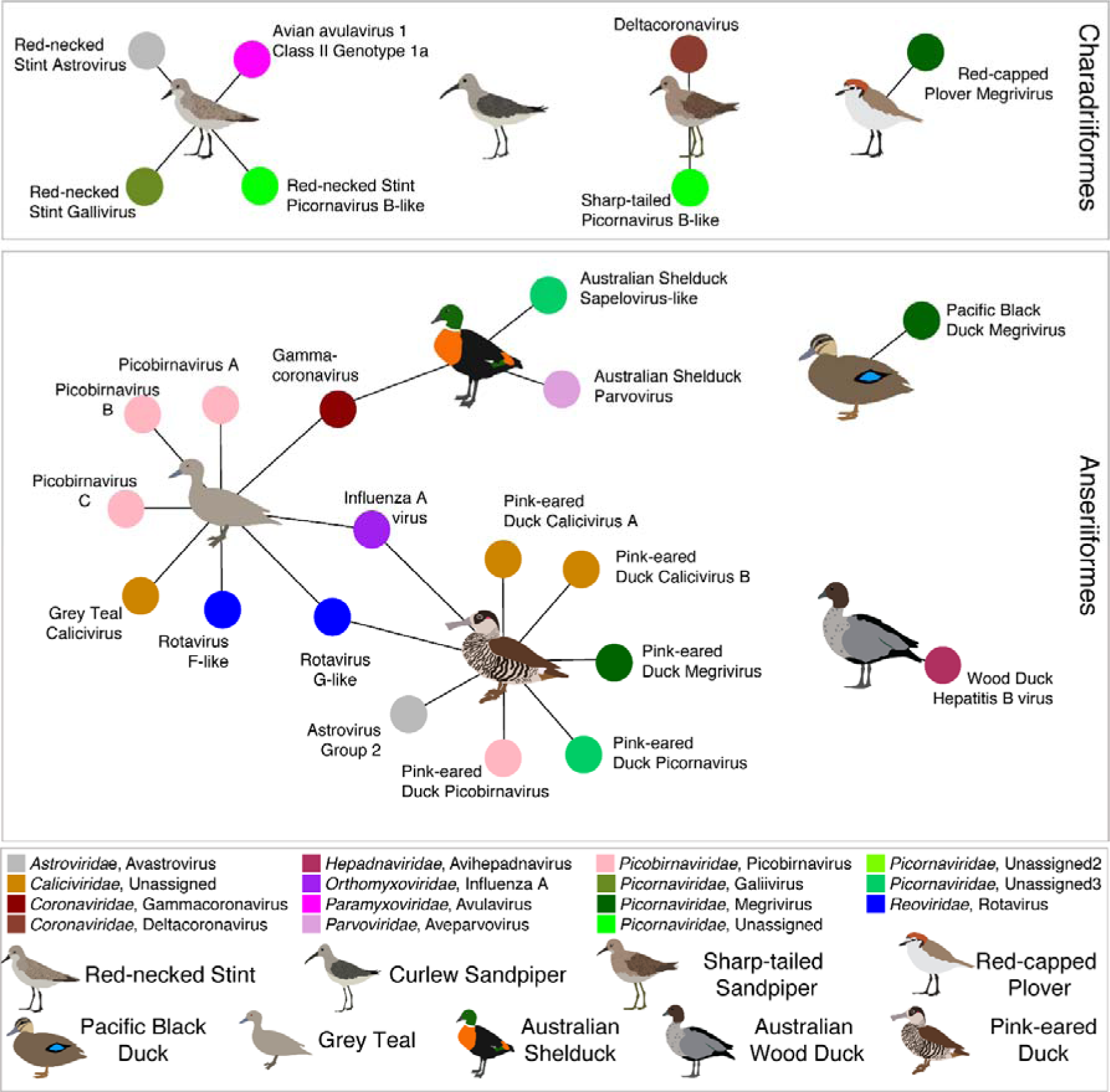
Bipartite network illustrating the species for which complete viral genomes were found in each library. Each library is represented as a central node, with a pictogram of the avian species, surrounded by each viral species. Where no complete viral genomes were revealed, the pictogram is shown with no viruses. Where two libraries share a virus species the networks between the two libraries are linked. Virus colour corresponds to virus taxonomy. A list of viruses from each library is presented in Table S3, and phylogenetic trees for each virus family and species can be found in (Fig 3-6, Figs S2-S14).

#### Novel ssRNA viruses

Two novel avastroviruses were identified. Red-necked Stint avastrovirus likely comprises a new virus “Group” as it falls basal to the Group 1 and 2 viruses in our phylogenetic analysis (Fig S3). Although no full genome Group 3 viruses exist, phylogenetic analysis of a short region of the RdRp demonstrated that this virus does not belong to Group 3 avastroviruses (Fig S4). Analysis of this short RdRp region also suggested that Red-necked Stint avastrovirus is sister to a virus detected in Swedish Mallards, indicating that this new group may be globally distributed (Fig S4). A Group 1 Avastrovirus, Pink-eared Duck astrovirus was also identified, and was a sister to Turkey astrovirus 2 (Fig S3, S4).

Our study further expanded an unassigned avian specific genus of the *Caliciviridae* with the identification of three new species, two from Pink-eared Ducks and one from Grey Teal. These viruses form a clade comprising Goose calicivirus, Turkey calicivirus and a calicivirus previously identified in Red-necked Avocets from Australia (*Recurvirostra novaehollandiae*) (Fig 3).

**Figure 3.**
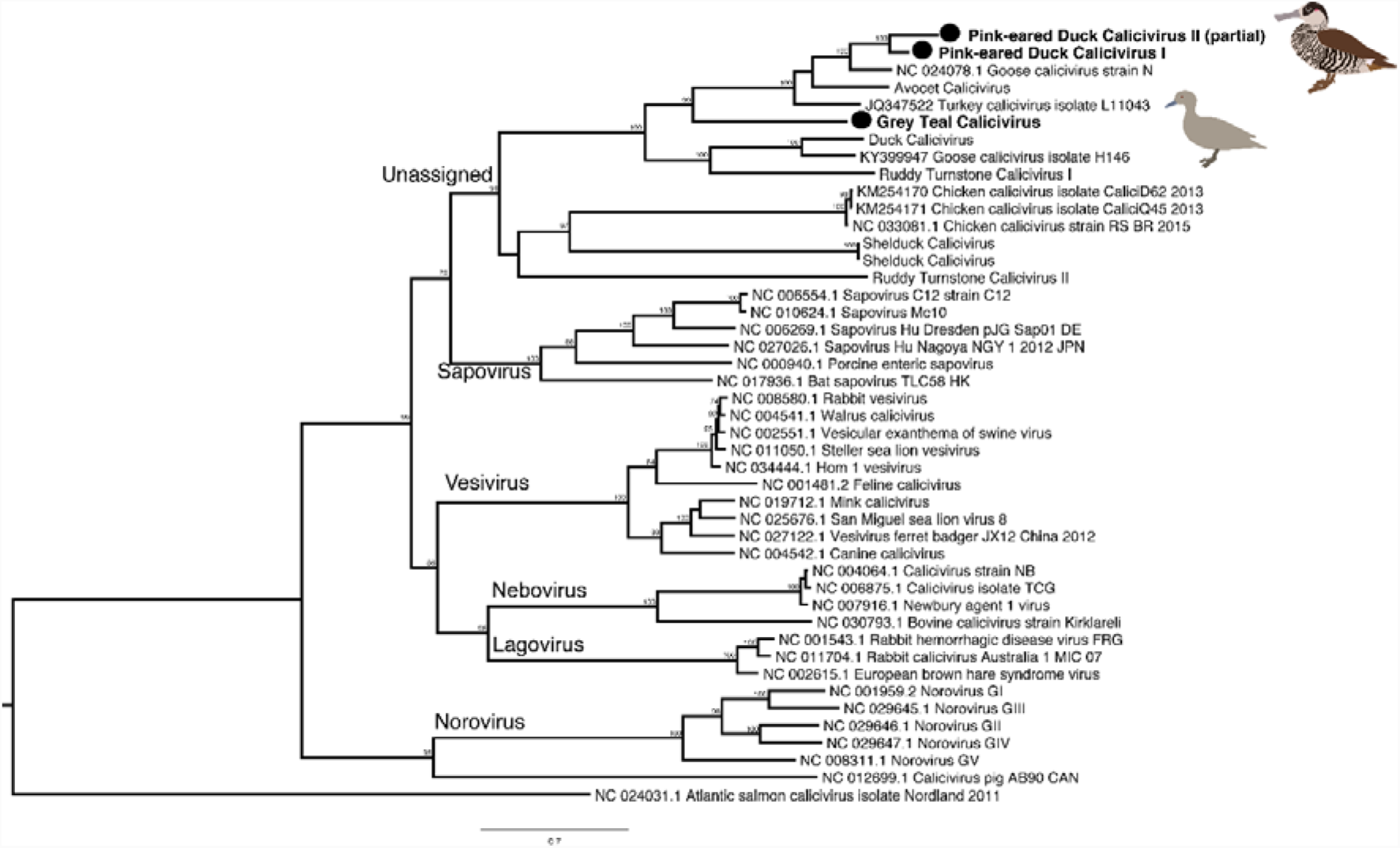
Phylogenetic tree of the virus polyprotein, including the RdRp, of representatives of the vertebrate RNA virus family the *Caliciviridae*. Viruses identified in this study are denoted with a filled circle and in bold. The most divergent calicivirus, Atlantic Salmon calicivirus, was used as outgroup to root the tree. Bootstrap values >70% are shown for key nodes. The scale bar represents the number of amino acid substitutions per site.

Members of the *Picornaviridae* were commonplace and genomes or partial genomes were identified in almost every library sequenced (Fig 4, Fig S5). Further, two different species of Picornaviruses were detected in the Red-necked Stint (Red-necked Stint Gallivirus and Red-necked Stint Picornavirus B-like) and Pink-eared Duck (Pink-eared Duck Megrivirus and Pink-eared Duck Picornavirus) libraries. To date, Galliviruses have only been isolated from Galliformes, so it was unexpected to identify a virus that was sister to this genus in the Red-necked Stint, although long branch lengths involved may indicate a novel viral genus. A virus similar to Avian sapeloviruses was identified in an Australian Shelduck (Fig 4, Fig S5). Three different Megriviruses were also identified, two from Anseriformes and one from Charadriiformes. Finally, a number of picornaviruses from unassigned genera similar to Pigeon picornavirus B were identified in Charadriiformes (Fig 4, Fig S5). These form a clade with a number of picornaviruses previously detected in Red-necked Avocets from Australia. Two host libraries had more than one picornavirus species (Red-necked Stint and Pink-eared Duck) (Fig 2).

**Figure 4.**
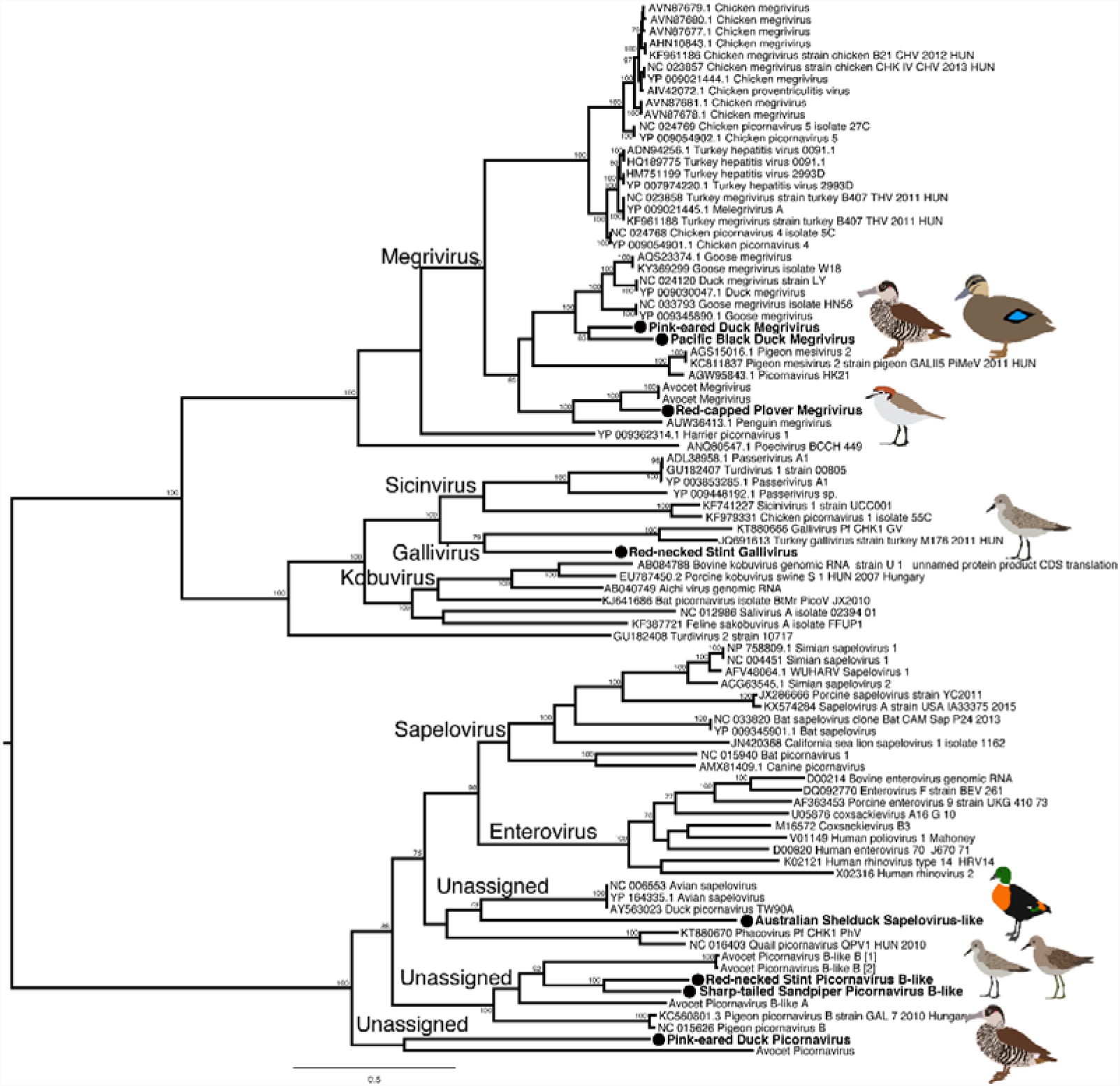
Phylogeny of the virus polyprotein, containing the RdRp, of selected members of the *Picornaviridae*. An expanded tree containing reference viruses for all main avian and mammalian genera is presented in Figure S5. The tree was midpoint rooted for clarity only. Viruses described in this study are marked in bold, adjacent to a filled circle. Bootstrap values >70% are shown for key nodes. The scale bar indicates the number of amino acid substitutions per site.

In addition to a previously described coronavirus, we identified a potentially novel species of deltacoronavirus. Specifically, sharp-tailed Sandpiper deltacoronavirus was most closely related to deltaviruses in wild birds from the United Arab Emirates and gulls from Europe (Fig S11), although the limited number of deltacoronaviruses sequences available inhibits a detailed analysis of its geographic range.

#### Novel dsRNA viruses

Three picobirnavirus species from two waterfowl species were newly identified here. Grey Teal picobirnavirus X and Pink-eared Duck picobirnavirus are members of a broad clade closely related to picobirnaviruses sampled in a number of species including Turkeys (from which only very short sequences are available and hence not analysed here (Fig 5). Grey Teal picobirnavirus Y falls into a divergent clade largely comprised of human and porcine picobirnaviruses (and no turkey picobirnaviruses), potentially representing an interesting host-switching event (Fig 5). However, due to limited sampling in wild birds, this virus could be related to other, currently unsampled, avian picorbirnaviruses. Two novel rotavirus species were also revealed from these host species. Indeed, Grey Teals and Pink-eared ducks shared a rotavirus species, distantly related to rotavirus G. This virus is one of three shared viruses in our entire dataset. Grey Teals also carried a second rotavirus species, distantly related to rotavirus F (Fig S7).

**Figure 5.**
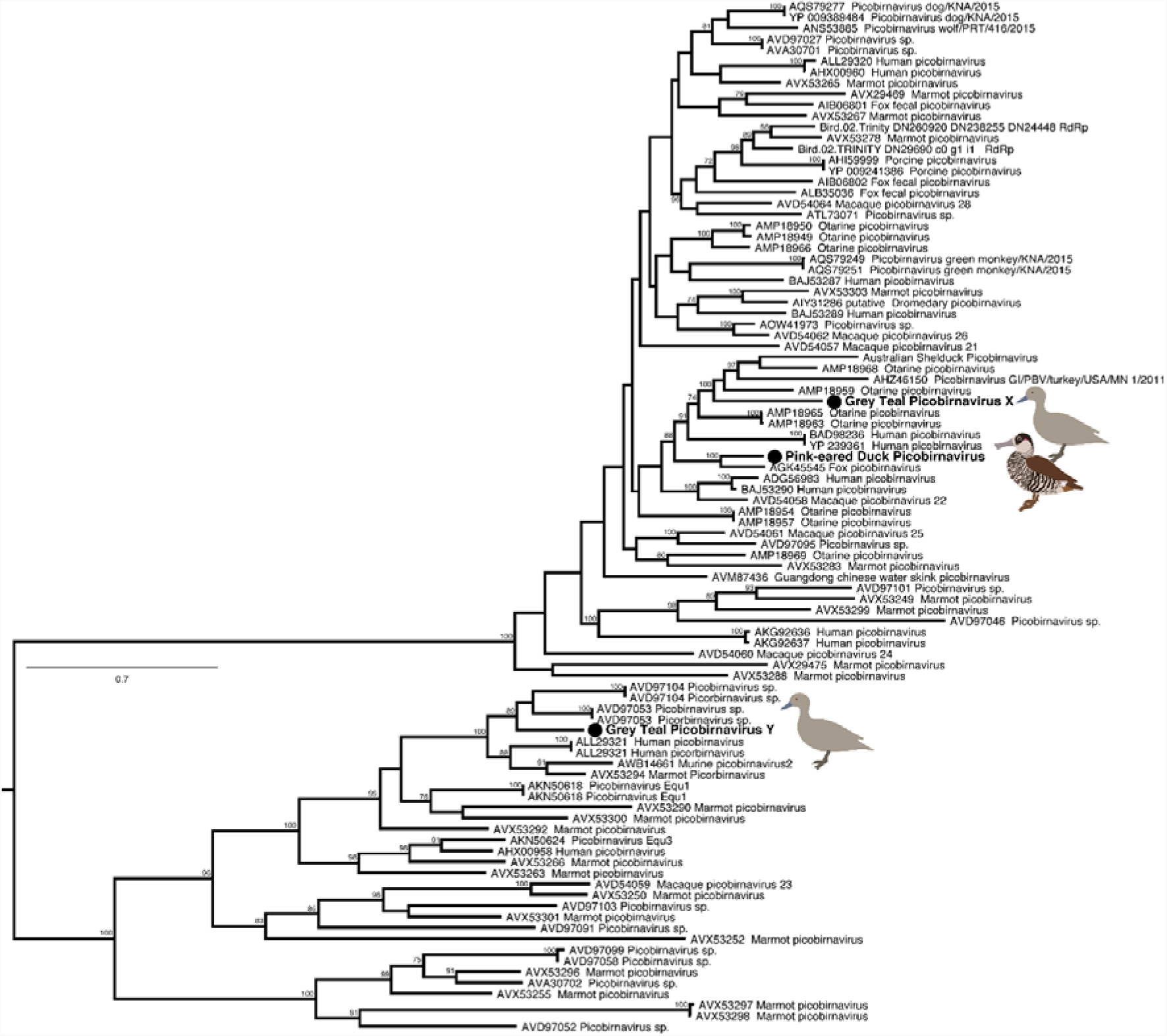
Phylogeny of virus segment 2, containing the RdRp, of the *Picobirnaviridae*. The tree was midpoint rooted for clarity only. Viruses described in this study are marked in bold, adjacent to a filled circle. Bootstrap values >70% are shown for key nodes. The scale bar indicates the number of amino acid substitutions per site.

#### Novel DNA viruses

These data also provided evidence for the presence of a novel single-stranded DNA virus and retro-transcribing DNA virus. An Australian Shelduck parvovirus (ssDNA) was revealed in the Grey Teal (Grey Teal Parvovirus) that belongs to the highly divergent genus Chapparvovirus of the *Parvoviridae* (Fig 6). Exogenous hepadnaviruses (retro-transcribing DNA viruses) from waterfowl are host specific, and the novel Wood Duck Hepatitis B virus identified here is most closely related to Shelgoose Hepatitis B virus, Duck Hepatitis B virus and Snow Goose Hepatitis Virus (Fig S8).

**Figure 6.**
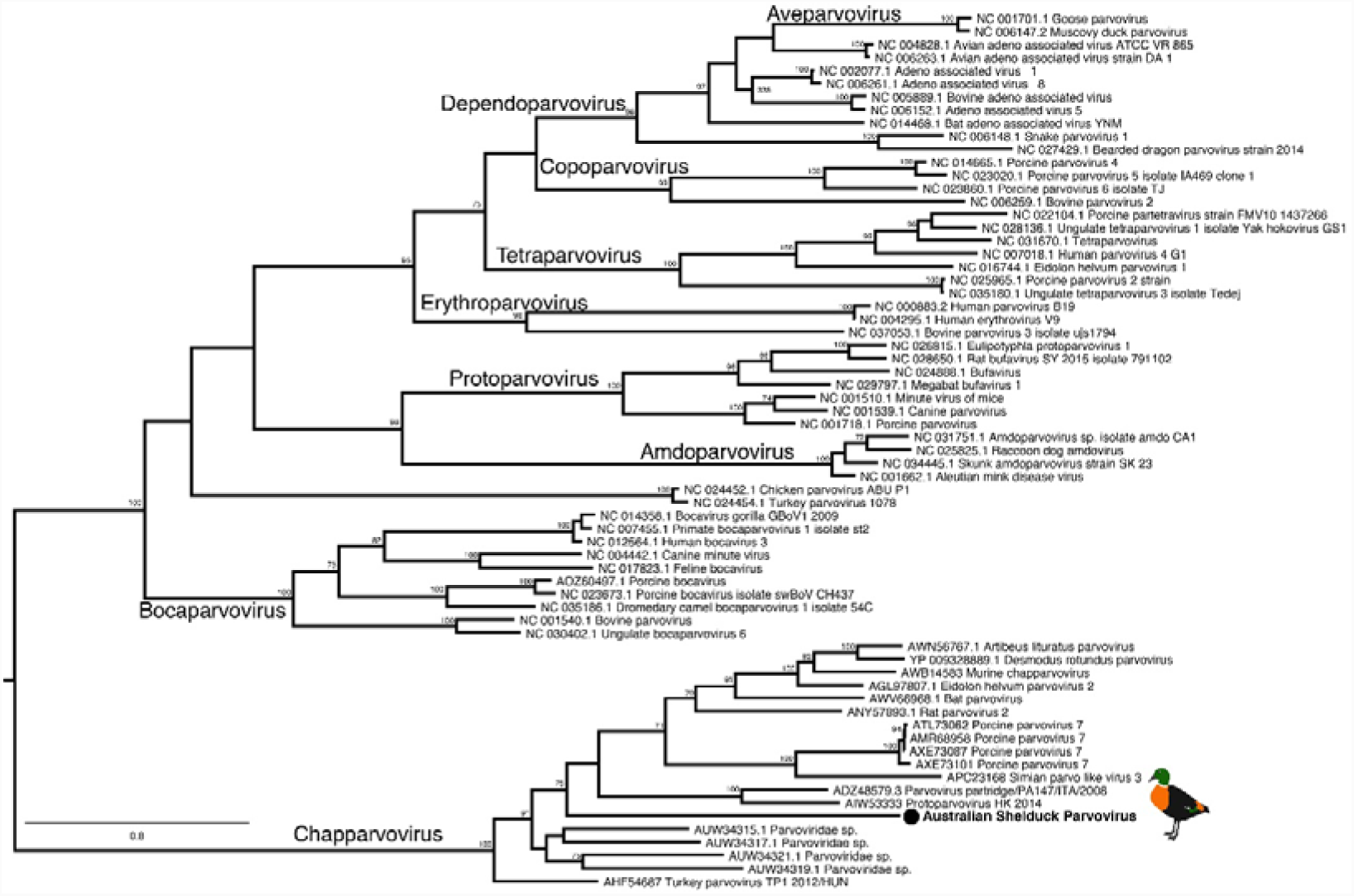
Phylogenetic tree of the NS protein of representative *Parvovirinae* (ssDNA). The sequence generated in this study are indicated by a filled circle and are shown in bold. The tree was midpoint rooted for clarity only. Bootstrap values >70% are shown for key nodes. The scale bar indicates the number of amino acid substitutions per site.

### Previously described avian RNA viruses

Given their frequency in avian populations as described in numerous surveillance schemes, we anticipated finding IAV, Avian avulavirus type 1 (formerly avian paramyxovirus type 1), and members of the *Coronaviridae* in some of the species sampled. Not only did we detect these viruses, but IAV and avian gammacoronavirus were shared across three different waterfowl libraries (Fig S9-S10). Phylogenetic analysis of a partial RdRp revealed that the avian gammacoronavirus identified here were most closely related to those already found in Australian wild birds (Fig S10).

We identified two subtypes of influenza A virus – H9N1 and H3N1 (Fig S12, S13) – in Grey Teal and Pink-eared Duck, respectively. Both H9N1 and H3N1 are rarely detected subtype combinations in large waterfowl surveillance schemes (47,48). Segments of these two viruses generally fell into the geographically isolated “Eurasian” clade, with the exception of the NP segment that fell within the “North American” clade, thereby demonstrating intercontinental reassortment in Australian waterfowl viruses (Fig S12, S13). Finally, although avian avulavirus Type 1 Class II Genotype 1b are frequently isolated from wild birds globally, we detected a Class II Genotype 1a virus infrequently isolated in wild birds (Fig S14). This genotype has been previously isolated from Australian chickens, although the F gene cleavage site (GRQGR*L) indicates this virus is of the low pathogenic phenotype.

### Host heterogeneity and connectivity of avian viromes

There was a large variation in the abundance of avian viral reads across the libraries. The highest abundance of avian viruses was in Red-necked Stint and Sharp-tailed Sandpiper, with 0.08% of reads in both cases; the lowest viral abundance was also in this avian order (Curlew Sandpiper, 0.00018% of reads). Of the Anseriformes, Grey Teal, Australian Shelduck and Pink-eared Duck had high abundance (0.051%, 0.035%, and 0.047% of reads, respectively) while Wood Duck and Pacific Black Duck had very low abundance, albeit only one order of magnitude lower (0.0024% and 0.0019%, respectively) (Table S1).

As with abundance, there was marked heterogeneity in alpha diversity (i.e. the diversity of viruses in each library) within the Anseriformes and Charadriiformes. In contrast to abundance, alpha diversity was higher in Anseriform than Charadriiform libraries at the viral family, genus, and species levels, although this difference was not statistically significant (Fig S15-S17). The lack of significance is potentially due to the surprisingly high alpha diversity in Red-necked Stint compared to other Charadriiformes, and the low alpha diversity in Pacific Black Ducks and Australian Wood Ducks in the Anseriformes (Fig 7A, Fig S15-S18). Hence, despite sampling multispecies flocks, there can be a large variation in virome structure across species and potentially individuals.

**Figure 7.**
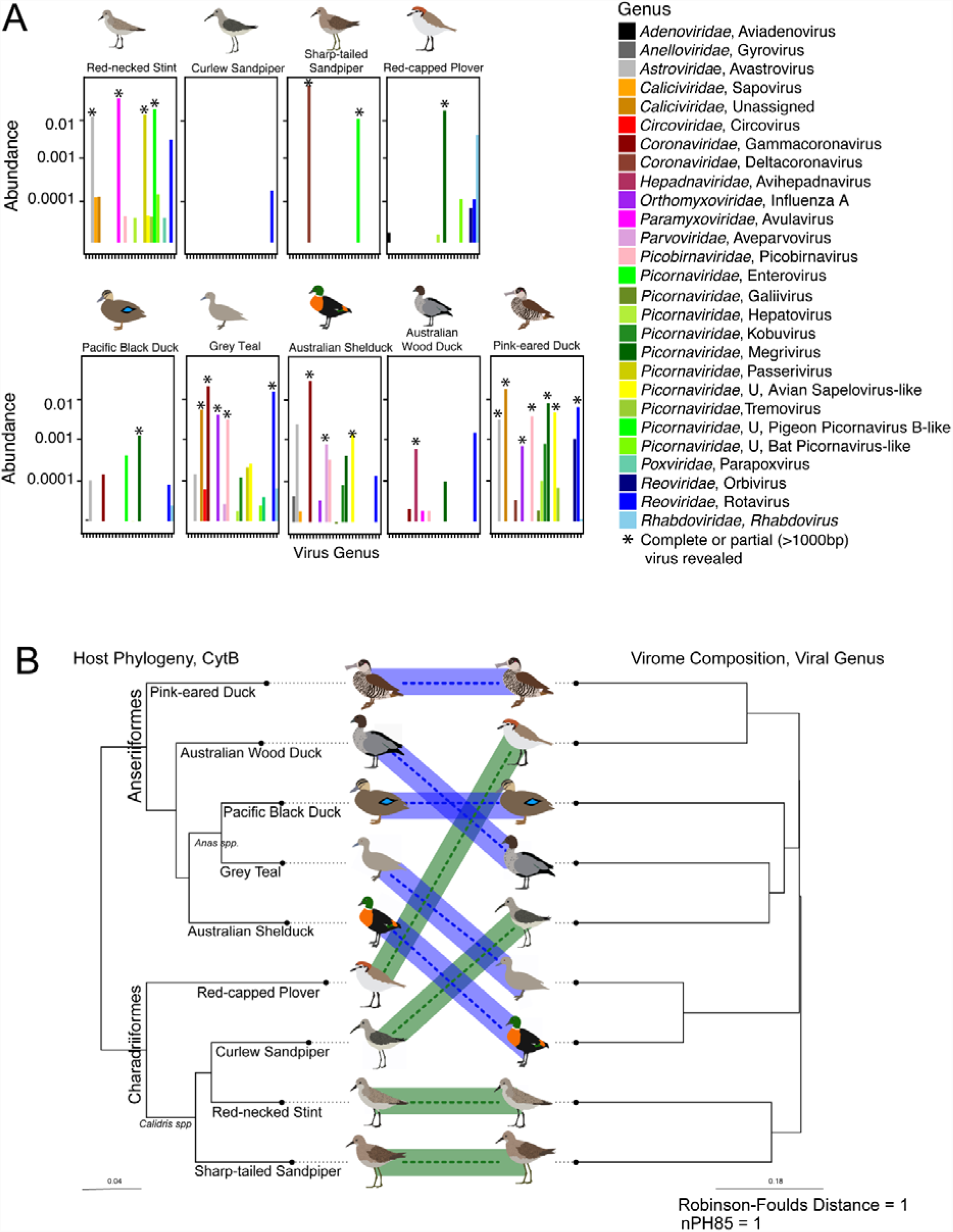
Heterogeneity and lack of taxonomic structure in avian viromes. (A) Abundance of avian viral genera identities in each library. Libraries are arranged taxonomically, with cladograms illustrating host species phylogenetic relationships within the Charadriiformes and Anseriformes. The taxonomic identification presented is that of the top blastx hit of all avian viral contigs. Asterisks indicate cases in which at least one complete or partial (>1000bp) virus was obtained. Alpha diversity metrics are presented in Fig S17. (B) Co-phylogeny demonstrating the discordance between host taxonomic relationship and virome composition. Host (phylogenetic) taxonomic relationship was inferred using the mitochondrial cytochrome B gene. Virome composition dendogram generated by clustering of bray-curtis dissimilarity matrix. The relationship between host taxonomy and virome composition was tested using two discordance metrics: Robinson-Foulds and nPH85, where 1 is discordance and 0 is agreement.

There was also substantial variation in the viral genera and species composition within each library (Fig 7A, Fig S15-S18). Members of the *Picornaviridae, Caliciviridae* and *Reoviridae* (genus *Rotavirus*) were ubiquitous and full genomes or short contigs were found in almost every library, often at high abundance (Fig 7A, Fig S14, S16). In addition to picornaviruses and rotaviruses, the Red-necked Stint had a highly abundant astrovirus (0.012%) and avulavirus (0.035%), while Sharp-tailed Sandpipers had a highly abundant deltacoronavirus (0.073%) that were not detected in other Charadriiform libraries (Fig 7A, S17). More viral families, genera and species were shared amongst the Anseriformes, particularly between Grey Teal, Pink-eared Duck and Australian Shelduck. Specifically, Grey Teal and Australian Shelduck shed avian gammacoronavirus at high abundance (0.021% and 0.029%, respectively) and Grey Teal and Pink-eared Duck shed IAV, although at lower abundances (0.0041% and 0.000728%, respectively) (Fig 7A, Fig S17). Overall, there were no significant trends towards differential abundance of viral families between the Charadriiformes and Anseriformes (Fig S19-S21). Viruses from the *Astroviridae, Calciviridae, Coronaviridae, Picobirnaviridae, Picornaviridae, Reoviridae* and *Rhabdoviridae* were found in both Anseriformes and Charadriiformes, albeit with different abundance patterns (Fig S19-S21). Multiple genera of picornaviruses were detected, but similarly without significant differences: most families and genera were detected in both avian orders (Fig S19-S21).

Using a non-metric ordination method (NMDS) we found no clustering of viral family or viral genus by host order with respect to virus abundance and diversity between libraries (i.e. beta-diversity) (Fig S22). Similarly, there was no statistically significant clustering of viral families (R^2^ = 0.13, df = 8, p = 0.267) or genera (R^2^ = 0.157, df = 8, p = 0.173) in the Charadriiformes compared to Anseriformes, suggesting that despite taxonomic differences waterbirds share numerous viral genera and families. We utilized a co-phylogenetic approach to better determine whether this lack of clustering was associated with host phylogeny. Accordingly, a phylogram of beta-diversity was not congruent with host phylogeny at the viral family, genus or species levels (Fig 7B, Fig S23-S24). Hence, evolutionary relationships among hosts may not play a major role in structuring viromes. For example, closely related sister species (the *Anas* ducks Grey Teal and Pacific Black Ducks, or *Calidris* sandpipers Sharp-tailed Sandpipers, Red-necked Stint and Curlew Sandpipers) do not possess viromes that are more similar to each other than to those of more distantly related species within the same avian order. Rather, the phylogram of beta diversity has clusters with a mix of Anseriform and Charadriiform libraries, indicating connectivity of viral families and genera across species of both avian orders (Fig 7B, Fig S23-S24). Similarly, there was also no statistically significant clustering by feeding mechanism in the Anseriformes (R^2^=0.56, df=4, p=0.2) or migratory propensity in the Charadriiformes (R^2^=0.43, df=3, p=0.25) (Fig S22) suggesting that host ecology may also play a limited role in shaping virome composition at the levels of viral family and genus.

Assessing comparative virome structure at the viral family and genera level is critical in demonstrating core viral families and genera in waterbirds. Species level analysis, albeit limited to viruses in which we were able to assemble >1000bp, is a more accurate measure of connectivity and heterogeneity of avian viromes. First, while there was no marked division in virome composition at the level of viral family or genera between the Anseriformes and the Charadriiformes, there was such a distinction at the level of viral species. Specifically, no viral species were shared between the Anseriformes and Charadriiformes (Fig 3), although the Anseriformes and Charadriiformes were sampled at different time points and at different locations. Within the Anseriformes, three viruses (influenza A virus, avian gammacoronavirus, and duck rotavirus G-like) were shared between three libraries: Grey Teal, Pink-eared Duck and Australian Shelduck (Fig 3, Fig S23). These shared viruses were especially common in the viromes of the Grey Teal (80% of avian viral reads), Australian Shelduck (82% of avian viral reads), and a small proportion of Pink-eared Duck avian viral reads (17%). The two *Anas* ducks (Grey Teal and Pacific Black Duck), the most closely related Anseriformes, did not share any viral species; surprisingly, the virome of the Pacific-Black Duck was different from the three connected host species. Further, Grey Teal and Pink-eared Ducks, the most taxonomically distinct waterfowl, shared two viral species, demonstrating the limited impact of host phylogeny (Fig S23). These viruses were also shared across ecological niches (dabbling ducks and filter feeding ducks), suggesting that co-occurrence was potentially responsible for their spread.

Within the Anseriformes we tested for the effect of virus-virus interactions on alpha diversity, specifically whether the presence of IAV had an effect on virome abundance or composition. Despite a low abundance of IAV in the libraries, there was a trend towards a higher alpha diversity of viromes in Anseriform species positive for IAV (Grey Teal, Pink-eared Duck; Fig S25) at the viral family, genus and species levels, although this was not statistically significant due to small sample size. Furthermore, IAV positive libraries were adjacent to each on the NMDS plots, although this clustering was not statistically significant (Viral Family R^2^=0.14, df=8, p=0.307, Viral Genera, R^2^=0.14, df=8, p=0.224). This trend did not hold for avian gammacoronavirus in the Grey Teal and Australian Shelduck libraries (Fig S26).

Finally, the phylogenetic analysis did not reveal a clear host-taxonomic gradient in viral species relationships. However, within the megriviruses (*Picornaviridae*), there appear to be large clades that may reflect avian order, with the viruses identified in the Anseriformes and Charadriiformes falling into two different clades. Furthermore, viruses from wild Anseriformes fall as sister taxa to previously described duck and goose megriviruses (Fig S6). In addition, this and our previous study (25) identified a number of picornaviruses from an unassigned genus only found in Charadriiformes, such that it might similarly represent a virus genus that is specific to a particular host order.

## Discussion

We identified 27 novel and previously described viral species from nine waterbirds falling into two avian orders. Anseriformes and Charadriiformes are important reservoirs for the best described avian virus, IAV, but are also central to the epidemiology of other multi-host viruses such as avian coronavirus and avian avulavirus type 1 (10, 11, 24, 49-53). As such, these avian hosts are excellent model species for understanding the determinants of virome composition. Indeed, we detected all these previously described low pathogenic avian viruses in our sample set, and coronaviruses and influenza A viruses were shared across different Anseriform species. We also genomically described 24 novel viral species belonging to 10 viral families, including both RNA and DNA viruses. The largest number and diversity of viruses belonged to the *Picornaviridae*, although a number of rotaviruses and caliciviruses were also described.

Overall, the avian viruses identified in this study were most closely related to other avian viruses, or in genera containing avian viruses. The exception is Grey Teal picorbirnavirus Y that occupies a clade dominated by viruses from human and porcine hosts. Whether this represents a host switch, or is due to lack of sampling in other hosts, will likely be revealed in additional metatranscriptomic studies. The Shelduck parvovirus described here is of particular interest as it is a member of the genus *Chapparvovirus*. Metagenomic analyses have recently identified members of this genus in a large number of vertebrates (54), and are known agents of severe disease (55).

Beyond viral discovery, our study revealed no predictable clustering of viromes according to host taxonomy in either the Anseriformes and/or Charadriiformes. Given the data on IAV, we might expect see differences in virome structure due to a number of host factors (11), including differences in biology/physiology. For example, different host species have different cell receptors which in turn results in different cell and tissue tropisms and patterns of viral attachment (56). Further, following infection, different species have differences in long-term immune memory (21,22). However, we saw no clear distinction between the viromes of Anseriformes or Charadriiformes based on host taxonomy, suggesting these host factors are not central to virome structuring. For example, within the Charadriiformes, the closely related *Calidris* sandpipers (Scolopacidae) did not have similar viromes and did not cluster as a group independently from Red-capped Plover, a member of a different avian family (Charadridae). Alternatively, it is possible that aspects of host ecology, such as foraging ecology, may be more important in shaping virome composition than host taxonomy (a proxy for physiology). Specifically, differences in ecology may generate differences in virus exposure across closely related hosts (11,57,58). The five Anseriform species studied here utilize three different feeding ecologies - dabbling, grazing and filter feeding - while the four Charadriiform species have different bill lengths and forage in different layers of sediment (12). Notably, however, there was no clear difference in virome composition as a result of ecology.

Central to our study was considering virome structure in the context of a multi-host and multi-virus model of virus-host interactions. Accordingly, the data generated here revealed large-scale heterogeneity (abundance and alpha diversity) in virome composition at the levels of virus family, genus and even species. Despite this heterogeneity, there was also some connectivity between viromes at the levels of virus family, genus and even species: some viral families and genera were ubiquitous in almost all avian libraries, including members of the *Picornaviridae* and *Reoviridae*. More striking was the connectivity between three avian species (Grey Teal, Australian Shelduck, and Pink-eared Duck) at the level of viral species: these hosts shared Influenza A virus, gammacoronavirus, and Duck G-like rotavirus. As these Anseriformes were sampled in the same temporal and spatial frames, it is not unexpected that there was connectivity despite differences in ecology and taxonomy. However, at the level of viral species there was no connectivity between the Anseriform and Charadriiform libraries, although this may be due to the physiological differences noted above or, more simply, that the ducks and wader viruses were sampled at different times and places. Despite the lack of connectivity between the Anseriformes and Charadriiformes at a viral species level, avian avulavirus 1 and deltacoronavirus detected in Red-necked Stint and Sharp-tailed Sandpiper, respectively, have been previously described in Anseriformes (10,51). This connectivity between viromes is likely facilitated by the association of these birds to water. Specifically, viruses such as IAV are thought to be primarily transmitted by the faecal-oral route, in which viruses contaminate water through the faeces and birds ingest the viruses while feeding or preening (15). Such water-borne transmission is critical to the dynamics of infection in bird communities (59). Furthermore, aquatic habitats seemingly support a higher risk of infection as compared to terrestrial habitats (58,60). In support of this was the observation of lower viral diversity and abundance in the grazing Australian Wood duck which has a more terrestrial dietary strategy compared to the other Anseriform species.

In sum, viral families and genera appeared to be readily shared among hosts, suggesting that waterbirds are key hosts for these families and genera. More importantly, our results indicate that avian viromes are largely comprised of what appear to be multi-host generalist viruses (here, influenza A virus, avian coronavirus, avian avulavirus type 1, duck rotavirus D-like) along with potential host-specific specialist viruses, which likely play a role in driving both heterogeneity and connectivity. While we found no evidence for viral species shared across avian orders, known multi-host virus species were revealed in both avian orders. Cases of clear host specificity were rare, but we speculate that Wood Duck Hepatitis B virus is likely host specific given high host specificity in this viral family (61). In addition, the clade level structuring of Megriviruses (*Picornaviridae*), and previous report of an identical Megrivirus species found in the same avian species in very different locations (25), similarly suggests some level of host specificity in this viral genus. Viral discovery efforts are imperative to better understanding factors that shape virome structure and the scope of host specificity in the avian reservoir. Importantly, we believe it is also imperative to consider multi-host, multi-virus systems in virus ecology.

## Acknowledgements

The sampling was supported by NIAID (HHSN266200700010C), ARC discovery grants (DP130101935 and DP160102146). The Melbourne WHO Collaborating Centre for Reference and Research on Influenza is supported by the Australian Department of Health. ECH is funded by an ARC Australian Laureate Fellowship (FL170100022). We thank the Victorian Wader Study Group the logistic support from Melbourne Water, and duck hunters who kindly sampled duck carcasses. We also thank Simeon Lisovski and Marta Ferenczi for their support

## References

1. Ostfeld RS,Keesing F. Effects of Host Diversity on Infectious Disease. Annu Rev Ecol Evol S. 2012;43:157–82.

2. Keesing F, Holt RD, Ostfeld RS. Effects of species diversity on disease risk. Ecol Lett. 2006;9(4):485–98.

3. Johnson PT, Preston DL, Hoverman JT, LaFonte BE. Host and parasite diversity jointly control disease risk in complex communities. Proc Natl Acad Sci U S A. 2013;110(42):16916–21.

4. LoGiudice K, Ostfeld RS, Schmidt KA, Keesing F. The ecology of infectious disease: Effects of host diversity and community composition on Lyme disease risk. Proc Natl Acad Sci U S A. 2003;100(2):567–71.

5. Milholland MT, Castro-Arellano I, Suzan G, Garcia-Pena GE, Lee TE, Rohde RE, et al. Global Diversity and Distribution of Hantaviruses and Their Hosts. EcoHealth. 2018;15(1):163–208.

6. Haydon DT, Cleaveland S, Taylor LH, Laurenson MK. Identifying reservoirs of infection: A conceptual and practical challenge. Emerg Infect Dis. 2002;8(12):1468–73.

7. Altizer S, Nunn CL, Thrall PH, Gittleman JL, Antonovics J, Cunningham AA, et al. Social organization and parasite risk in mammals: Integrating theory and empirical studies. Annual Review of Ecology Evolution and Systematics. 2003;34:517–47.

8. Streicker DG, Fenton A, Pedersen AB. Differential sources of host species heterogeneity influence the transmission and control of multihost parasites. Ecology Letters. 2013;16(8):975–84.

9. Geoghegan JL, Holmes EC. Predicting virus emergence amid evolutionary noise. Open Biol. 2017;7(10).

10. Wille M, Avril A, Tolf C, Schager A, Larsson S, Borg O, et al. Temporal dynamics, diversity, and interplay in three components of the viriodiversity of a Mallard population: Influenza A virus, avian paramyxovirus and avian coronavirus. Infect Genet Evol. 2015;29:129–37.

11. van Dijk JGB, Verhagen JH, Wille M, Waldenström J. Host and virus ecology as determinants of influenza A virus transmission in wild birds. Curr Opin Virol. 2018;28:26–36.

12. Dann P. Foraging behaviour and diets of red-necked stints and curlew sandpipers in south-eastern Australia. Wildlife Research. 2000;27(1):61–8.

13. Wikramaratna PS, Pybus OG, Gupta S. Contact between bird species of different lifespans can promote the emergence of highly pathogenic avian influenza strains. Proc Natl Acad Sci U S A. 2014;111(29):10767–72.

14. Ren HG, Jin Y, Hu MD, Zhou J, Song T, Huang ZS, et al. Ecological dynamics of influenza A viruses: cross-species transmission and global migration. Sci Rep-Uk. 2016;6.

15. Olsen B, Munster VJ, Wallensten A, Waldenström J, Osterhaus ADME, Fouchier RAM. Global patterns of influenza A virus in wild birds. Science. 2006;312:384–8.

16. Munster VJ, Baas C, Lexmond P, Waldenström J, Wallensten A, Fransson T, et al. Spatial, temporal, and species variation in prevalence of influenza A viruses in wild migratory birds. PloS Pathog. 2007;3:e61. doi:10.1371/journal.ppat.0030061.

17. Maxted AM, Luttrell MP, Goekjian VH, Brown JD, Niles LJ, Dey AD, et al. Avian influenza virus infection dynamics in shorebird hosts. J Wildl Dis. 2012;48(2):322–34.

18. Bröjer C, Ågren EO, Uhlhorn H, Bernodt K, Mörner T, Jansson DS, et al. Pathology of natural highly pathogenic avian influenza H5N1 infection in wild tufted ducks (*Aythya fuligula*). J Vet Diagn Invest. 2009;21:579–87.

19. Pantin-Jackwood MJ, Costa-Hurtado M, Shepherd E, DeJesus E, Smith D, Spackman E, et al. Pathogenicity and Transmission of H5 and H7 Highly Pathogenic Avian Influenza Viruses in Mallards. J Virol. 2016;90(21):9967–82.

20. Kim J-K, Negovetich NJ, Forrest HL, Webster RG. Ducks: the “trojan horses” of H5N1 influenza. Influenza and Other Respiratory Viruses. 2009;3:121–8.

21. Magor KE. Immunoglobulin genetics and antibody responses to influenza in ducks. Dev Comp Immunol. 2011;35:1008–16.

22. Hill SC, Manvell RJ, Schulenburg B, Shell W, Wikramaratna PS, Perrins C, et al. Antibody responses to avian influenza viruses in wild birds broaden with age. Proceedings Biological sciences. 2016;283(1845):doi:10.1098/rspb.2016.159.

23. Ferenczi M. Avian influenza virus dynamics in Australian wild birds. PhD Thesis: Deakin University; 2016.

24. Ferenczi M, Beckmann C, Warner S, Loyn R, O’Riley K, Wang X, et al. Avian influenza infection dynamics under variable climatic conditions, viral prevalence is rainfall driven in waterfowl from temperate, south-east Australia. Vet Res. 2016;47:23.

25. Wille M, Eden JS, Shi M, Klaassen M, Hurt AC, Holmes EC. Virus-virus interactions and host ecology are associated with RNA virome structure in wild birds. Mol Ecol. 2018: doi:10.1111/mec.14918.

26. Shi M, Lin XD, Tian JH, Chen LJ, Chen X, Li CX, et al. Redefining the invertebrate RNA virosphere. Nature. 2016;540(7634):539–43.

27. Shi M, Lin XD, Chen X, Tian JH, Chen LJ, Li K, et al. The evolutionary history of vertebrate RNA viruses. Nature. 2018;556(7700):197–202.

28. Langmead B, Salzberg SL. Fast gapped-read alignment with Bowtie 2. Nat Methods. 2012;9(4):357–U54.

29. Katoh K, Standley DM. MAFFT multiple sequence alignment software version 7: improvements in performance and usability. Mol Biol Evol. 2013;30(4):772–80.

30. Capella-Gutierrez S, Silla-Martinez JM, Gabaldon T. trimAl: a tool for automated alignment trimming in large-scale phylogenetic analyses. Bioinformatics. 2009;25(15):1972–3.

31. Guindon S, Dufayard JF, Lefort V, Anisimova M, Hordijk W, Gascuel O. New algorithms and methods to estimate maximum-likelihood phylogenies: assessing the performance of PhyML 3.0. Systematic biology. 2010;59(3):307–21.

32. Lagkouvardos I, Fischer S, Kumar N, Clavel T. Rhea: a transparent and modular R pipeline for microbial profiling based on 16S rRNA gene amplicons. Peerj. 2017;5:e2836. doi:10.7717/peerj.2836.

33. Oksanen J, Kindt R, Legendre P, O’Hara B, Stevens MHH, Oksanen MJ, et al. The vegan package. Community Ecology Package. 2007;10:631–7.

34. McMurdie PJ, Holmes S. phyloseq: An R package for reproducible interactive analysis and graphics of microbiome census data. PloS ONE. 2013;8(4):e61217. doi:10.1371/journal.pone.0061217.

35. Brooks AW, Kohl KD, Brucker RM, van Opstal EJ, Bordenstein SR. Phylosymbiosis: Relationships and Functional Effects of Microbial Communities across Host Evolutionary History. PloS Biology. 2016;14(11).

36. Robinson DF, Foulds LR. Comparison of Phylogenetic Trees. Math Biosci. 1981;53(1-2):131–47.

37. Schliep KP. phangorn: phylogenetic analysis in R. Bioinformatics. 2011;27(4):592–3.

38. Penny D, Hendy MD. The Use of Tree Comparison Metrics. Syst Zool. 1985;34(1):75–82.

39. Geoghegan JL, Duchene S, Holmes EC. Comparative analysis estimates the relative frequencies of co-divergence and cross-species transmission within viral families. PloS Pathog. 2017;13(2).

40. Paradis E, Claude J, Strimmer K. APE: Analyses of Phylogenetics and Evolution in R language. Bioinformatics. 2004;20(2):289–90.

41. Barth JMI, Matschiner M, Robertson BC. Phylogenetic Position and Subspecies Divergence of the Endangered New Zealand Dotterel (Charadrius obscurus). PloS One. 2013;8(10).

42. Baker AJ, Pereira SL, Paton TA. Phylogenetic relationships and divergence times of Charadriiformes genera: multigene evidence for the Cretaceous origin of at least 14 clades of shorebirds. Biol Lett-Uk. 2007;3(2):205–9.

43. Gibson R, Baker A. Multiple gene sequences resolve phylogenetic relationships in the shorebird suborder Scolopaci (Aves: Charadriiformes). Mol Phylogenet Evol. 2012;64(1):66–72.

44. Sun ZL, Pan T, Hu CC, Sun L, Ding HW, Wang H, et al. Rapid and recent diversification patterns in Anseriformes birds: Inferred from molecular phylogeny and diversification analyses. PloS One. 2017;12(9).

45. Revell LJ. phytools: an R package for phylogenetic comparative biology (and other things). Methods Ecol Evol. 2012;3(2):217–23.

46. Love MI, Huber W, Anders S. Moderated estimation of fold change and dispersion for RNA-seq data with DESeq2. Genome Biology. 2014;15(12).

47. Latorre-Margalef N, Tolf C, Grosbois V, Avril A, Bengtsson D, Wille M, et al. Long-term variation in influenza A virus prevalence and subtype diversity in a migratory Mallards in Northern Europe. Proceedings Biological sciences. 2014;281:doi:10.1098/rspb.2014.0098

48. Wilcox BR, Knutsen GA, Berdeen J, Goekjian VH, Poulson R, Goyal S, et al. Influenza A viruses in ducks in Northwestern Minnesota: fine scale spatial and temporal variation in prevalence and subtype diversity. PloS ONE. 2011;6:e24010. doi:10.1371/journal.pone.0024010.

49. Wille M, Lindqvist K, Muradrasoli S, Olsen B, Jarhult JD. Urbanization and the dynamics of RNA viruses in Mallards (*Anas platyrhynchos*). Infect Genet Evol. 2017;51:89–97.

50. Wille M, Muradrasoli S, Nilsson A, Jarhult JD. High prevalence and putative lineage maintenance of avian coronaviruses in Scandinavian waterfowl. PloS ONE. 2016;11(3):e0150198. doi:10.1371/journal.pone.

51. Chamings A, Nelson TM, Vibin J, Wille M, Klaassen M, Alexandersen S. Detection and characterisation of coronaviruses in migratory and non-migratory Australian wild birds. Sci Rep-Uk. 2018;8.

52. Hoque MA, Burgess GW, Cheam AL, Skerratt LF. Epidemiology of avian influenza in wild aquatic birds in a biosecurity hotspot, North Queensland, Australia. Preventive veterinary medicine. 2015;118(1):169–81.

53. Hoque MA, Burgess GW, Karo-Karo D, Cheam AL, Skerratt LF. Monitoring of wild birds for Newcastle disease virus in north Queensland, Australia. Preventive veterinary medicine. 2012;103(1):49–62.

54. de Souza WM, Romeiro MF, Fumagalli MJ, Modha S, de Araujo J, Queiroz LH, et al. Chapparvoviruses occur in at least three vertebrate classes and have a broad biogeographic distribution. J Gen Virol. 2017;98(2):225–9.

55. Roediger B, Lee Q, Tikoo S, Cobbin JCA, Henderson JM, Jormakka M, et al. An Atypical Parvovirus Drives Chronic Tubulointerstitial Nephropathy and Kidney Fibrosis. Cell. 2018;175(2):530–43 e24.

56. Jourdain E, van Riel D, Munster V, Kuiken T, Waldenström J, Olsen B, et al. The pattern of influenza virus attachment varies among wild bird species. PloS ONE. 2011;6(9):doi:10.1371/journal.pone.0024155.

57. Hall SR, Sivars-Becker L, Becker C, Duffy MA, Tessier AJ, Caceres CE. Eating yourself sick: transmission of disease as a function of foraging ecology. Ecology Letters. 2007;10(3):207–18.

58. Lozano GA. Optimal Foraging Theory - a Possible Role for Parasites. Oikos. 1991;60(3):391–5.

59. Roche B, Lebarbenchon C, Gauthier-Clerc M, Chang CM, Thomas F, Renaud F, et al. Water-borne transmission drives avian influenza dynamics in wild birds: the case of the 2005-2006 epidemics in the Camargue area. Infect Genet Evol. 2009;9(5):800–5.

60. Hoye BJ, Fouchier RAM, Klaassen M. Host behaviour and physiology underpin individual variation in avian influenza virus infection in migratory Bewick’s Swans. Proceedings Biological sciences. 2012;279(1728):529–34.

61. Guo H, Mason WS, Aldrich CE, Saputelli JR, Miller DS, Jilbert AR, et al. Identification and characterization of avihepadnaviruses isolated from exotic anseriformes maintained in captivity.J Virol. 2005;79(5):2729–42.

